# Delineating the posterior bank of the feline auditory cortex based on neurofilament proteins expressing SMI-32

**DOI:** 10.64898/2026.07.28.741385

**Authors:** Austin Robertson, Jeffrey G. Mellott, Blake E. Butler

## Abstract

The feline auditory cortex is understood to consist of 13 distinct subregions with unique anatomical and functional properties. Differential patterns of SMI-32 immunoreactivity are commonly used to identify the borders between these subregions; however, the detailed description of areal differences that is commonly cited did not include descriptions of the patterns observed along the posterior ectosylvian gyrus. Thus, the current manuscript aims to provide a more complete data set that can used to delineate the dorsal, intermediate, and ventral divisions of the posterior ectosylvian gyrus (auditory cortical regions dPE, iPE, and vPE, respectively) based on SMI-32 reactivity using the same methods and measures. Taken together, the current data and those published previously allow for a standardized approach to identifying all 13 auditory cortical subregions in this essential model of auditory cortical structure and function.

## 1. Introduction

The accurate and comprehensive delineation of cortical subregions has been critical to the study of neuroanatomy and its relationship to function, pathology, and plasticity. Historically, borders between anatomically distinct brain areas have been defined based on differences in cytoarchitecture, myeloarchitecture, histochemistry, patterns of neuronal projections, or some combination of these measures. Often, this anatomical diversity gives rise to measurable functional differences in, for example, receptive field properties, patterns of functional connectivity, or behavioral outputs that further support delineation and characterization of individual subregions.

Developing these detailed relationships between cortical structure and function in a way that affords meaningful inference about the human cortex requires tractable animal models. With respect to the study of auditory processing, the cat has emerged as a powerful model due in part to the presence of a large and accessible auditory cortex containing regions with human homologues (Lomber and Malhotra, 2008). Within the feline auditory cortex (as in other species), the location/density of neuronal cell bodies and patterns of myelination can be used to visualize cortical laminae; however, the boundaries between adjacent regions are often less clear. Fortunately, the monoclonal antibody SMI-32 allows for visualization of differences in the distribution of neurofilament protein across feline auditory cortical regions that allows for the identification of areal borders (Mellott et al., 2010). Patterns of SMI-32 labelling have also been used to identify specific auditory cortical subregions across a variety of other species (e.g., Hof et al., 1992; Budinger et al., 2000; Lewis & Van Essen, 2000; Hassiotis et al., 2004; Ashwell et al., 2005; Wong & Kaas, 2009), as well as to delineate the feline visual cortex (Van der Gucht et al., 2001).

Patterns of SMI-32 labelling have been used to define regions of interest in studies of anatomical connectivity (e.g., Chabot et al, 2015; Butler et al 2016a; 2016b; 2017; 2018) and functional properties (e.g. Kok et al., 2015; Land et al., 2016; Merrikhi, 2022) of the regions comprising the feline auditory cortex. However, while the detailed analysis provided by Mellott and colleagues (2010) described patterns of labelling across ten cortical regions, that accounting did not describe SMI-32 reactivity in the dorsal, intermediate, and ventral fields of the posterior ectosylvian gyrus (dPE, iPE, and vPE, respectively) - regions commonly considered to comprise the posterior limit of auditory cortex (Lee and Winer, 2011; Hackett, 2011). As a result, researchers commonly need to combine patterns of SMI-32 reactivity with coarse delineation based on gyral/sulcal boundaries to identify the boundaries of the full complement of auditory cortical regions. This is problematic, as a comprehensive view of the role these posterior ectosylvian regions play in the auditory processing remains to be developed. Lee and Winer (2011) categorized the subregions of the posterior ectosylvian gyrus as putative multisensory areas based on the presence of thalamic projections arising from extralemniscal nuclei and dense patterns of corticocortical projections from visual and somatosensory areas. Indeed, their placement of these regions near the top of the proposed functional hierarchy in auditory cortex is supported by sparse patterns of myelination (Robertson et al., 2025) – a common feature of higher level, integrative sensory regions. Moreover, large but distinct projections to the claustrum originate from both iPE and dPE (Beneyto & Prieto, 2001), suggesting these areas may play a role in sensory awareness (Remedios et al., 2014) and mediating selective attention (Goll et al., 2015; Atlan et al., 2018). However, despite this body of anatomical evidence suggesting potentially crucial contributions to sound processing, very few studies have directly measured functional properties of these areas. As with any attempt to associate brain structure and function, these studies will require robust and repeatable ways to delineate cortical regions in order to validate that their functional measures (e.g., in-vivo electrophysiological or optical recordings) were confined to the intended location. To address this gap, the current work describes how patterns of SMI-32 reactivity vary across the posterior ectosylvian gyrus (i.e., between regions dPE, iPE, vPE). Specifically, we provide quantitative descriptions and photomicrographs that can be used in combination with the descriptions provided by Mellott and colleagues (2010) to provide a standardized method of delineating all 13 regions of the feline auditory cortex, and facilitate comprehensive study of the structure and function of auditory cortex in this important model of sensory system function.

## 2. Materials and Methods

Five adult domestic cats were included in these analyses. All procedures were conducted in accordance with the Canadian Council on Animal Care’s *Guide to the Care and Use of Experimental Animals* (Olfert et al. 1993), and were approved by the University of Western Ontario’s Animal Care Committee.

At the time of sacrifice, animals were premedicated with a combination of ketamine (4 mg/kg, i.m.) and dexdomitor (0.03 mg/kg, i.m.) and a catheter was inserted into the cephalic vein. Sodium pentobarbital (40 mg/kg, i.v.) was administered to induce general anesthesia, and heparin (1 mL: anticoagulant) and 1% sodium nitrite (1 mL: vasodilator) were administered intravenously. Each animal was then transcardially perfused with physiological saline (1 L), followed by 4% paraformaldehyde (2 L), and 10% sucrose (2 L). Following perfusion, brains were blocked in the coronal plane between Horsley-Clarke levels A18 and P2 (a region spanning the entirety of auditory cortex; Horsley & Clarke, 1908), removed, and immersed in a 30% sucrose cryoprotective solution for approximately 7 days. Brains were then sectioned in the coronal plane at a thickness of 60 um using a cryostat (Thermo-Fisher, Waltham, Mass). Every sixth section (360 um intervals) was processed using the monoclonal antibody SMI-32 to visualize patterns of neurofilament expression as described below, allowing for laminar and areal delineation of auditory cortical subregions.

### 2.1. SMI-32 Immunohistochemistry

SMI-32 is a monoclonal antibody that binds to non-phosphorylated epitopes on neurofilament proteins, allowing for the visualization of neuronal cell bodies, dendrites, and some thick axons in the central and peripheral nervous system of humans and other mammalian species (Mellot et al. 2010; Campbell and Morrison, 1989; Ang et al. 1991; Peichl, 1989; Straznicky et al. 1992; Han et al. 2021). Importantly, these epitopes are differentially expressed across regions and laminae of the auditory cortex, allowing them to be delineated (Mellott et al. 2010). To visualize patterns of SMI-32 reactivity, sections were rinsed with 0.1M physiological buffer (PB; 3 x 5 min), and endogenous peroxidase was blocked with 0.5% hydrogen peroxide in 70% ethanol (30 min). Sections were then rinsed with 0.1 M PB (4 x 5 min), incubated in 5% normal goat serum in 0.1M PB (45 min), and incubated in a mouse-anti-SMI-32 antibody (Vector Laboratories, Burlingame, CA, USA; 1/2000 in 2% NGS in 0.1M PB) overnight. The following day, sections were rinsed with 0.1 M PB (4 x 5 min) and incubated in biotinylated goat-anti-mouse IgG antibody (Vector Laboratories: 1/200 in 2% NGS in 0.1M PB) for 30 min. Sections were then rinsed with 0.1 M PB (3 x 10 min), and incubated in an avidin–biotin– horseradish peroxidase solution (Vectastain Elite ABC, Vector Laboratories) for 90 minutes.

Sections were then rinsed with 0.1 M PB (3 x 10 mins), and incubated with a diaminobenzene– nickel chromogen solution for 15 min. Finally, sections were rinsed with 0.1 M PB (3 x 5 min), mounted from 0.01 M PB on gelatinized slides, air-dried, cleared, and coverslipped.

### 2.2. Quantification of Somata and Dendrites

Mounted sections were examined using a Zeiss AxioImager M2 brightfield microscope equipped with a Lumina HR camera and StereoInvestigator software (MBF Bioscience, Inc., Williston, VT). Quantitative measurements were averaged across five sections from each animal for each region of interest (dPE, iPE, and vPE) as well as the dorsal zone of auditory cortex (DZ) to allow the estimates from those regions to be normalized to the estimates of Mellott and colleagues (2010). The boundaries of DZ were determined according to previous descriptions (Mellott et al., 2010). First approximations of the boundaries of dPE, iPE, and vPE were defined based on anatomical landmarks and the probabilistic atlas of the feline cortex (CATLAS; Stolzberg et al. 2017).

Sampling was performed by superimposing a counting frame at the centre of each region of interest that spanned the entirety of the cortical column, with a width of 500 μm for soma counts/measurement and 50 μm for dendrite counts. Within each counting frame, somata were counted, somatic diameter was measured, and dendrites were counted exhaustively using the meander scan function in Stereoinvestigator either using a 20x (soma) or 40x (dendrites) objective. Finally, counts and diameters were averaged across five sections to generate estimates for each animal, region of interest, and cortical layer.

### 2.3. Details of Normalization

While patterns of SMI-32 reactivity have become a standard approach for parcellating the cortex, variability in staining intensity is apparent between tissue samples processed asynchronously. Moreover, different observers may have different criteria for what constitutes a labelled soma or dendrite, even while using the same equipment or after having been given the same instructions. Thus, to ensure that the results of the current report can be compared directly to, and used alongside those of Mellott and colleagues (2010), data from the current study were normalized. Specifically, somatic and dendritic counts obtained from each layer of areas dPE, iPE, and vPE were divided by estimates obtained from the same layer of the dorsal zone of auditory cortex (DZ) within animal, and multiplied by the mean estimates for that layer in the DZ obtained by Mellott and colleagues (2010; see [1] for an example of how the soma count for a given area/layer was normalized). DZ was chosen as the basis of this normalization due to its highly identifiable pattern of SMI-32 reactivity across animals.

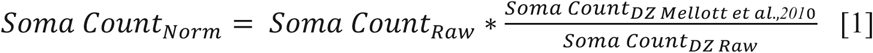

## 3. Results

### 3.1. SMI-32 Reactivity

Figure 1a shows a lateral view of the feline cortex, with auditory cortical subregions shaded and labeled. The dorsal, intermediate, and ventral fields of the posterior ectosylvian gyrus (dPE, iPE, and vPE) comprise the posterior limit of auditory cortex, occupying a large proportion of the tissue between the posterior ectosylvian sulcus and the posterior aspect of the suprasylvian sulcus. An example section showing SMI-32 reactivity across auditory cortical regions is shown in Figure 1b. In line with previous observations, SMI-32 labelling was greatest in layers III and V, with layers I and IV showing no immunoreactive neurons. Layers II and VI show weak-to-moderate labelling that varies by region. As expected, staining intensity correlates positively with total soma and dendrite counts. Moreover, there is a characteristic gradient in reactivity across the dorsoventral axis, with dorsal regions showing the strongest SMI-32 reactivity and ventral regions showing more sparse labelling (Fig. 1b). The mean raw soma counts, soma size estimates, and dendrite counts obtained for each region and layer are provided in Table 1. Mean soma size across region and layer is presented in Figure 2.

**Figure 1.**
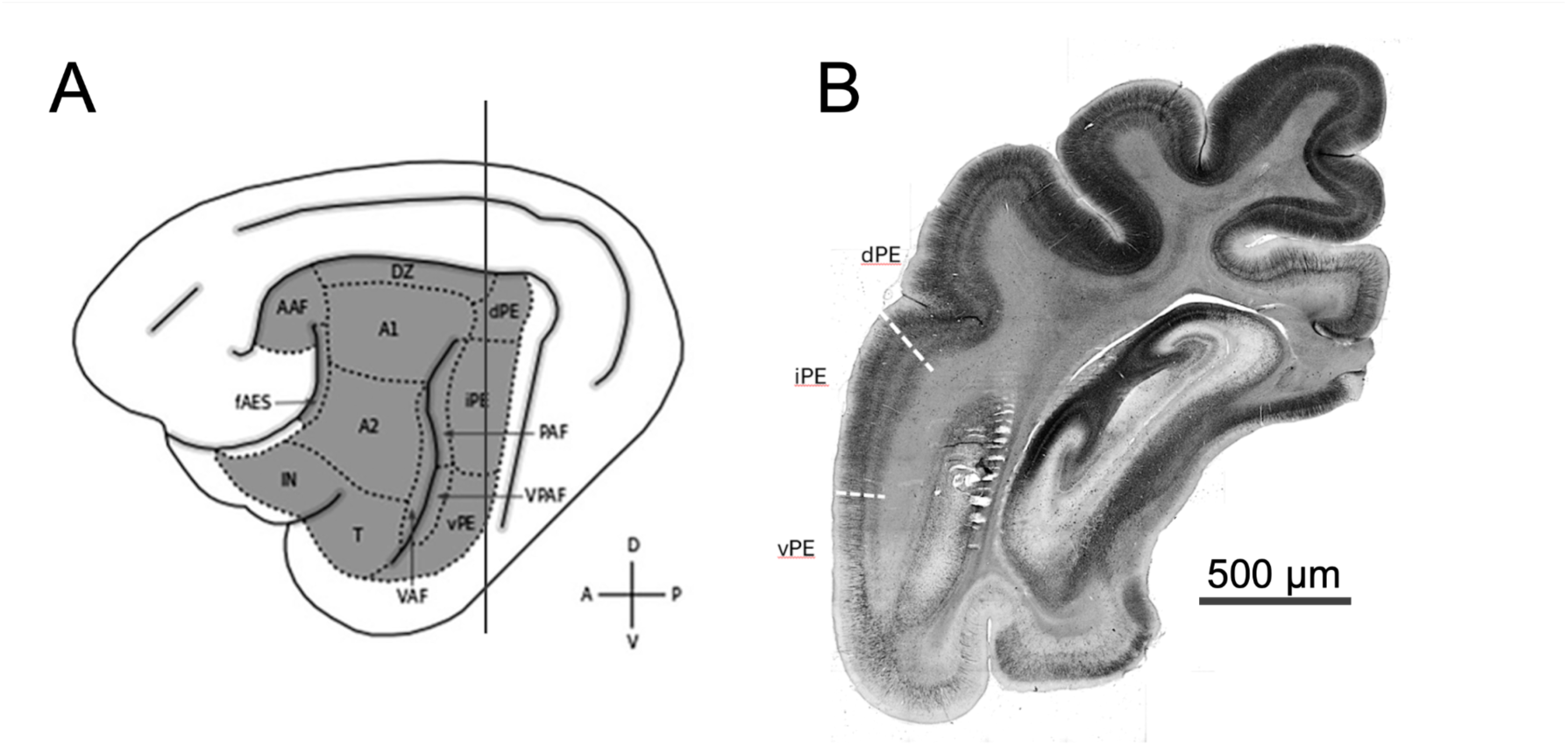
(A) Lateral view of the left cerebrum of the cat with the auditory cortex highlighted, and its constituent regions labelled. A line indicates the coronal position of the section pictured in B. (B) Macroscopic view of a coronal section with borders between dPE, iPE, and vPE estimated based on SMI-32 immunoreactivity. Even at low magnification, the differential staining of SMI-32 is apparent across regions, as is the dorsoventral gradient of immunoreactivity. AAF = anterior auditory field; DZ = dorsal zone of auditory cortex; A1 = primary auditory cortex; A2 = second auditory cortex; IN = insular area; T = temporal area; PAF = posterior auditory field; VPAF = ventral posterior auditory field; VAF = ventral auditory field; dPE/iPE/vPE = dorsal/intermediate/ventral aspects of the posterior ectosylvian gyrus.

**Figure 2.**
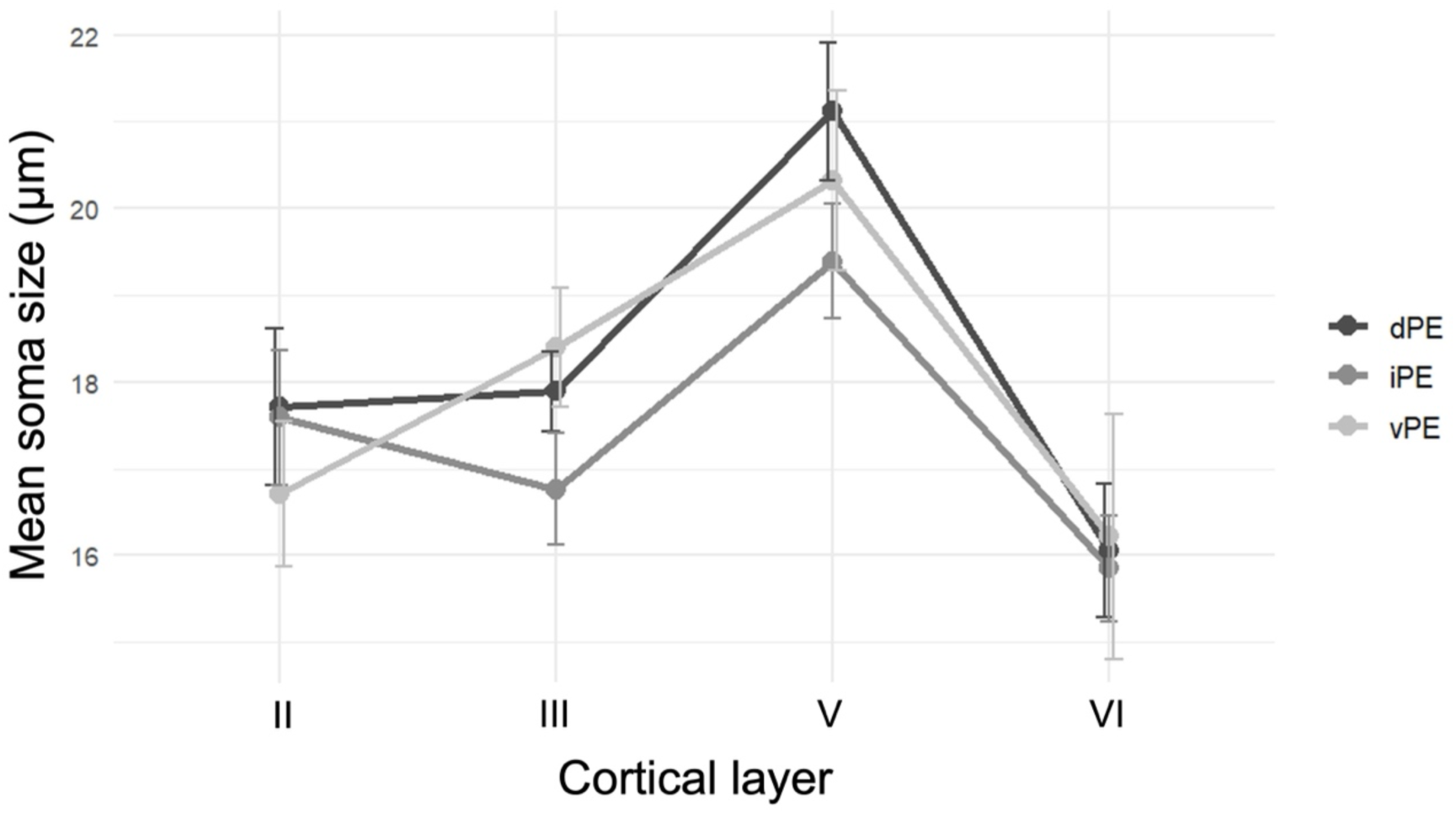
Mean soma size across regions and layers of the posterior ectosylvian gyrus. Error bars show the standard error of the mean. dPE/iPE/vPE = dorsal/intermediate/ventral aspects of the posterior ectosylvian gyrus.

**Table 1.**
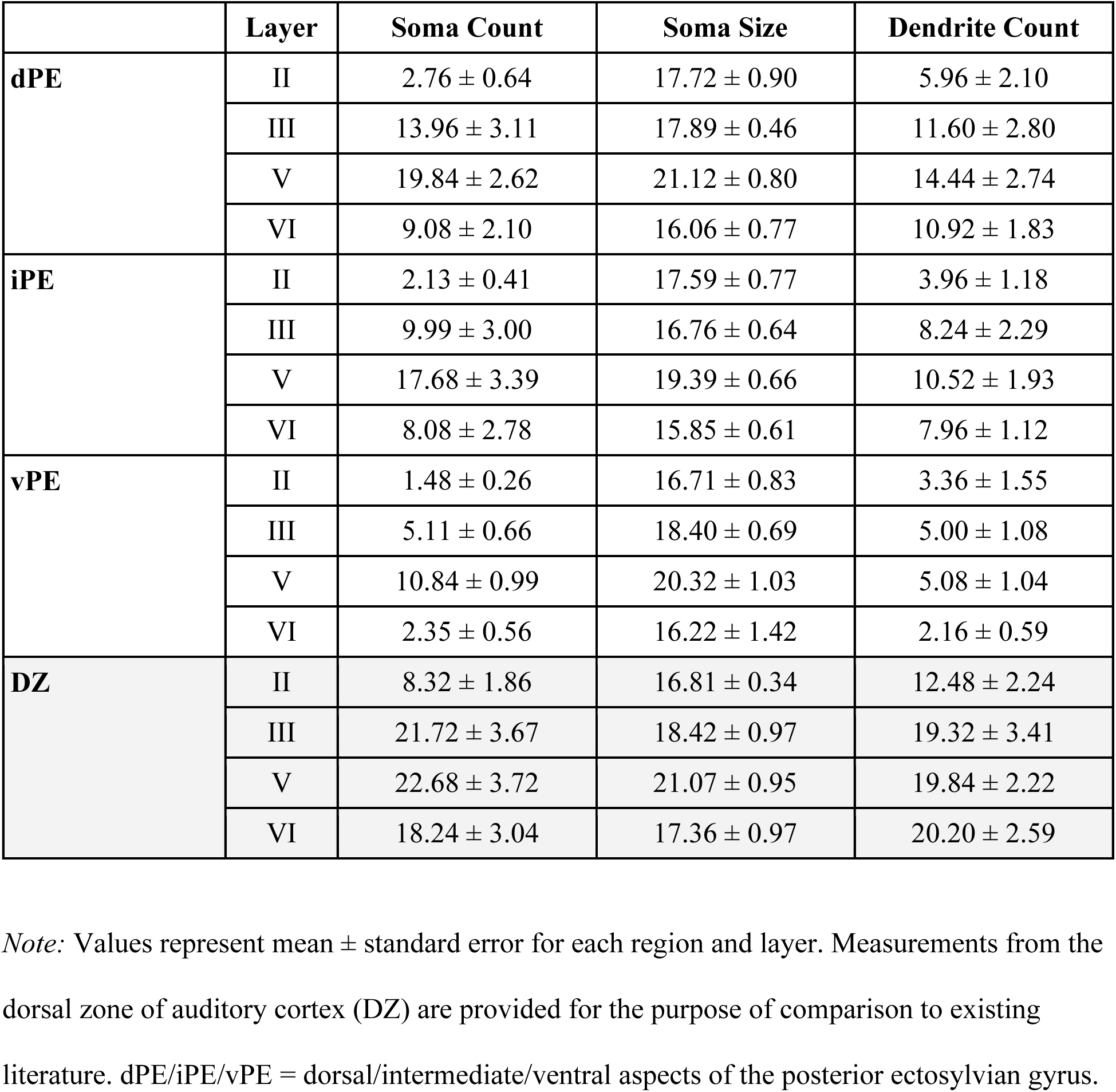
Raw soma counts, sizes, and dendrite counts across regions and layers of the posterior ectosylvian gyrus.

The intensity of SMI-32 labelling can vary across preparations in a way that reflects differences in experimental protocol rather than meaningful differences between animals (i.e., the reagents, antibody concentration, protocols, or perceptual thresholds for determining a cell has been labelled may differ across labs). Since one aim of the current study was to provide data that could be used in conjunction with those obtained from the other regions comprising the feline auditory cortex (Mellott et al., 2010), soma and dendrite counts were normalized such that they can be compared directly to previously obtained values (Table 2). Normalized mean soma and dendrite counts are presented in figures 3 and 4, respectively.

**Figure 3.**
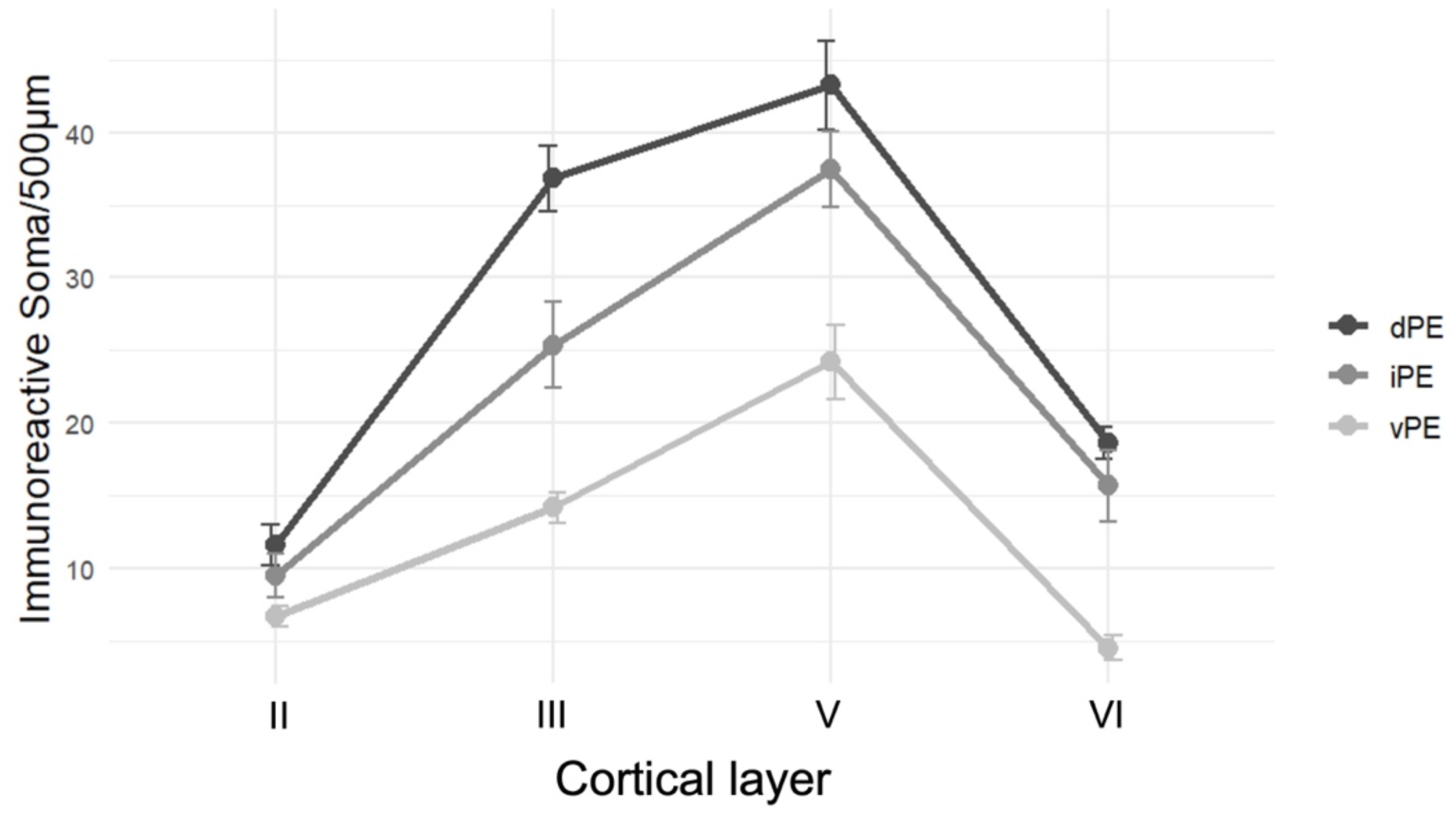
Mean normalized soma counts across regions and layers of the posterior ectosylvian gyrus. Error bars show the standard error of the mean. dPE/iPE/vPE = dorsal/intermediate/ventral aspects of the posterior ectosylvian gyrus.

**Table 2.** Soma and dendrite counts across regions and layers of the posterior ectosylvian gyrus.

|  | Layer | Soma Count | Dendrite Count |
| --- | --- | --- | --- |
| **dPE** | II | 11.56 ± 1.38 | 12.11 ± 1.11 |
|  | III | 36.82 ± 2.30 | 13.06 ± 2.76 |
|  | V | 43.26 ± 3.08 | 29.01 ± 3.66 |
|  | VI | 18.59 ± 1.12 | 30.20 ± 2.88 |
| **iPE** | II | 9.47 ± 1.51 | 8.78 ± 0.46 |
|  | III | 25.36 ± 2.94 | 8.86 ± 1.51 |
|  | V | 37.47 ± 2.57 | 20.90 ± 3.51 |
|  | VI | 15.65 ± 2.48 | 21.78 ± 1.84 |
| **vPE** | II | 6.68 ± 0.68 | 2.20 ± 0.32 |
|  | III | 14.14 ± 1.07 | 6.08 ± 2.10 |
|  | V | 24.21 ± 2.57 | 10.39 ± 1.49 |
|  | VI | 4.49 ± 0.84 | 13.03 ± 1.93 |

*Note:* Values represent mean ± standard error for each region and layer. Values obtained in the current study have been normalized as described in the text such that they are directly comparable to those obtained from by Mellott and colleagues (2010). dPE/iPE/vPE = dorsal/intermediate/ventral aspects of the posterior ectosylvian gyrus.

#### 3.1.1. Dorsal Posterior Ectosylvian Area

dPE occupies the dorsal aspect of the posterior ectosylvian gyrus. It has a heterogeneous staining pattern, with layers II and VI showing very little SMI-32 reactivity, and few labelled neurons or dendrites (Tables 1 & 2; Figures 3 & 4). Despite this relatively sparse labelling, layer VI appears somewhat thicker than adjacent regions (Figure 1b). Layer III has a moderate degree of immunoreactivity, with a substantial increase in somatic count (Tables 1 & 2; Figure 3) and size (Table 1; Figure 2), as well as a greater number of labelled dendrites than layer II (Tables 1 & 2; Figure 4). Finally, layer V shows strong SMI-32 reactivity, with more/larger somata (Table 1; Figures 2 & 3) and more labelled dendrites (Tables 1 & 2; Figure 4) than in any other layer.

**Figure 4.**
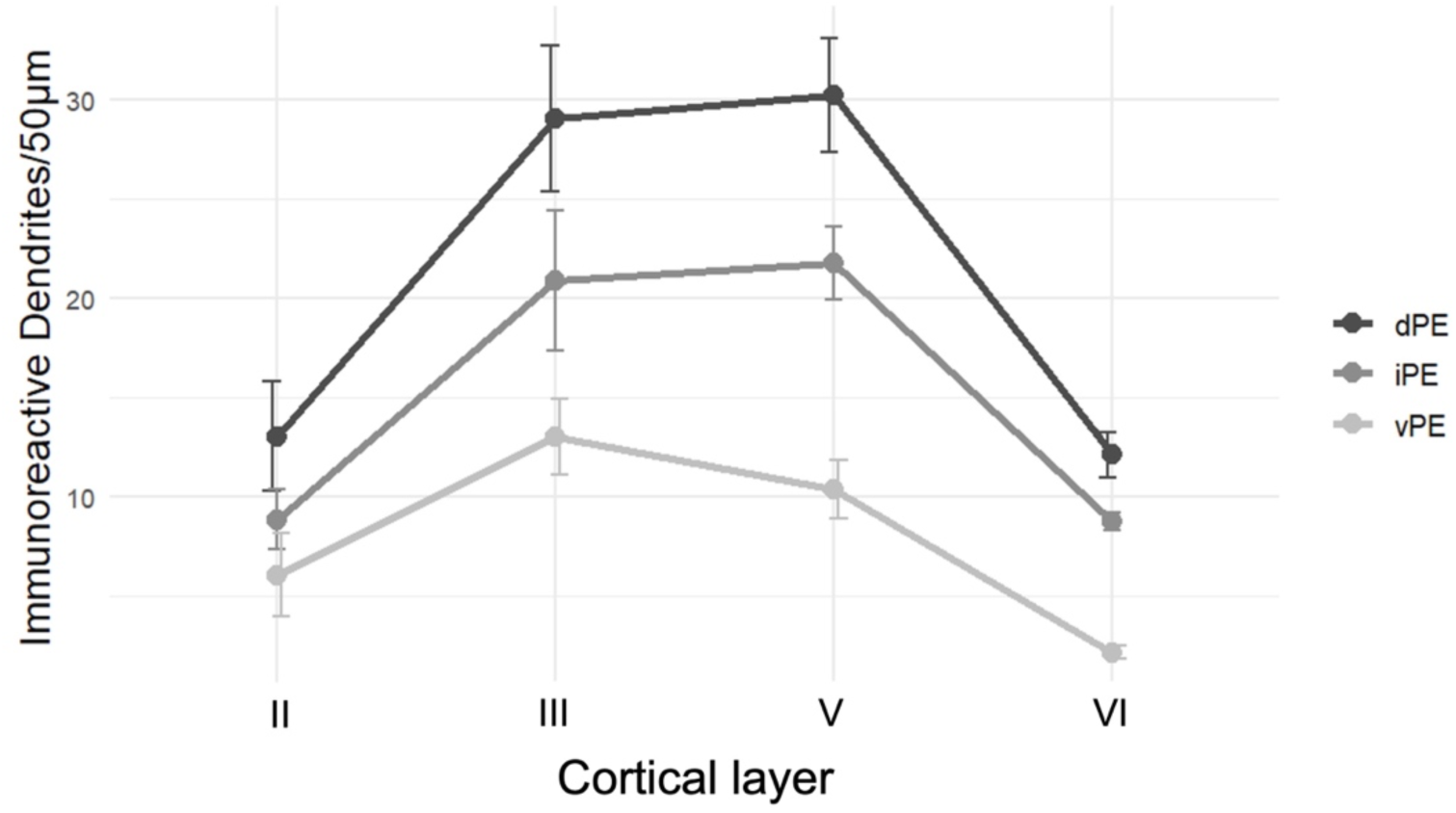
Mean normalized dendrite counts across regions and layers of the posterior ectosylvian gyrus. Error bars show the standard error of the mean. dPE/iPE/vPE = dorsal/intermediate/ventral aspects of the posterior ectosylvian gyrus.

As has been observed in other auditory cortical regions, no labelled somata are observed in layers I or IV. However, dendrites originating in layers II/III and V/VI travel through layers I/IV respectively. All layers are easily discernible at low magnification (Figure 1b).

#### 3.1.2. Intermediate Posterior Ectosylvian Area

iPE occupies the intermediate portion of the posterior ectosylvian gyrus, just ventral to dPE. It shows a similar general pattern of SMI-32 reactivity and structural features as dPE; however, iPE shows less reactivity across layers. Specifically, layer II has the fewest labelled somata and dendrites (Tables 1& 2; Figures 3 & 4), with some counting frames containing no labelled cells. Layers VI and III show slightly more reactivity, with layer V showing the greatest number of labelled soma/dendrites (Tables 1& 2; Figures 3 & 4). Generally, cell size and dendritic reactivity follow this same pattern across layers with slightly smaller soma (Table 1; Figure 2) and lower dendritic counts (Tables 1& 2; Figures 3 & 4) compared to those observed in dPE.

#### 3.1.3. Ventral Posterior Ectosylvian Area

vPE occupies the ventral portion of the posterior ectosylvian gyrus. It follows a similar pattern to the other posterior ectosylvian regions, with immunoreactivity, the number of labelled somata/dendrites (Tables 1& 2; Figures 3 & 4) and soma size (Table 1; Figure 2) being lowest in layers II and VI, followed by layer III, and finally layer V. However, substantially less SMI-32 reactivity is observed in vPE compared to both dPE and iPE, with many counting frames containing no labelled cells in layers II or VI. This sparse immunoreactivity makes the borders between vPE and adjacent auditory cortical regions readily identifiable (Figure 1b).

### 3.2. Boundary Discrimination and Delineation

Borders between adjacent cortical regions were identified based on differences in soma count/size and laminar patterns of SMI-32 reactivity, with gross anatomical features (e.g., sulcal/gyral landmarks) and patterns of myelination (Robertson et al., 2024) obtained from adjacent sections used as supporting evidence. Per the criteria established by Mellott and colleagues (2010), boundaries between regions lying along the posterior ectosylvian gyrus were deemed to be ‘clear-cut’, meaning the changes in SMI-32 reactivity appeared to be relatively abrupt (as opposed to ‘transitional’ borders where more subtle, gradual changes in reactivity are observed).

#### 3.2.1. dPE/iPE Border

The boundary between dPE and iPE (Figure 5a) can be distinguished at low magnification (although not as clearly as the border between iPE and vPE, as seen in Figure 5b).

**Figure 5.**
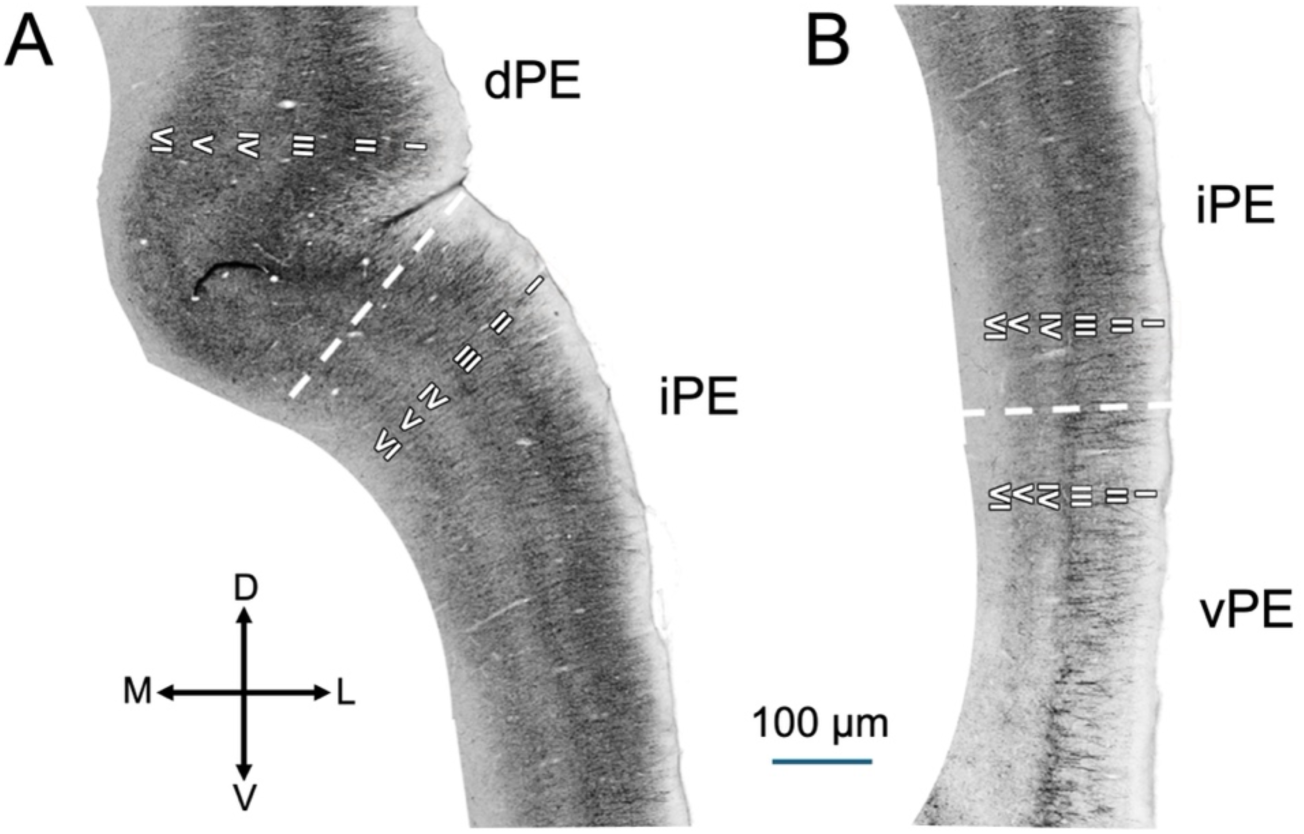
(A) The anatomical boundary between dPE and iPE is clear-cut, signaled by the relative differences in SMI-32 reactivity in layers III and V. In addition, dPE contains a greater number of dendrites originating in layer V and traveling through layer IV. (B) The anatomical boundary between iPE and vPE is clear-cut and distinguished by more robust reactivity in layer V of iPE relative to the sparse labelling in vPE. Cortical lamina are indicated in Roman numerals near the boundaries, denoted by dashed lines.

Identifying the boundary is facilitated by difference in the intensity of SMI-32 reactivity in layers III and V, which is greater in dPE than iPE, reflecting the greater number of stained somata in dPE (Tables 1 & 2; Figure 3), and their relatively larger size (Table 1; Figure 2). Furthermore, the two can be distinguished by the larger number of immunoreactive dendrites in layer IV of dPE (Tables 1 & 2; Figure 4), which originate in layer V and travel up through the cortical column.

#### 3.2.2. iPE/vPE border

The boundary between iPE and vPE (Figure 5b) is clear-cut, and is most easily identified based on differences in infragranular layers. While labelling in layers II and III is generally more intense in iPE than vPE, the reactivity is minimal in both regions, and sparsely-labelled sections can make delineation difficult. However, there is a robust difference in somatic labelling in layer V, where the moderately-labelled iPE gives way to the more sparsely labelled vPE (Tables 1 & 2; Figures 3 and 5b). In addition, the infragranular layers of the cortical column begin to widen near the border of the two regions.

## 4. Discussion

Accurate and standardized delineation of the distinct regions that comprise the cortex is central to defining hierarchical organization across the brain, and understanding how anatomical differences between regions might give rise to unique functional features. For the study of sensory perception, the cat has emerged as an important animal model from which many inferences about human perception have been drawn. Studies of the feline auditory cortex have identified 13 regions that can be distinguished based on their unique anatomical and/or functional properties. These delineations could be useful in identifying unique patterns of neuronal projections (e.g., Lee and Winer, 2008), or to confirm the location of recording electrodes (e.g. Mellott et al. 2010; Grieves et al. 2016; Stitt et al. 2018), recent studies have overwhelmingly turned to patterns of SMI-32 reactivity to visualize areal borders. As had been previously demonstrated in visual cortex (Van der Gucht, 2001), Mellott and colleagues (2010) demonstrated that ten of the 13 regions comprising the auditory cortex could be readily distinguished based on their laminar profiles of SMI-32 reactivity. The current study extends this work to include the 3 most posterior regions of feline auditory cortex - the dorsal, intermediate, and ventral divisions of the posterior ectosylvian gyrus. The ability to accurately delineate these regions based on cytoarchitectural features is essential to future efforts to better understand the unique functional roles of these subregions. This is particularly important given their purported role as high level, multisensory regions of the auditory pathway (Lee and Winer, 2011) that show robust connectivity outside the auditory cortex, and which may make meaningful contributions to processing and responding to complex sensory environments.

Here we provide estimates of soma counts, soma size, and dendrite counts across the cortical layers of these three regions. Moreover, we describe observable shifts in the patterns of SMI-32 reactivity that occur at the borders between adjacent regions, such that these areas can be delineated from one another. Finally, we provide normalized estimates of soma and dendrite counts such that the data from the current study can be used in concert with the estimates provided by Mellott and colleagues (2010), despite any potential differences in SMI-32 immunohistochemistry. Together, these two quantitative analyses of SMI-32 reactivity provide a comprehensive account that can be used to delineate the feline auditory cortex in its entirety and allow interpretation and integration of results in a common spatial context (Chon et al. 2019).

In addition to the differences reported between regions, we observed a clear dorsoventral gradient in SMI-32 reactivity, whereby the dorsal aspect of the posterior ectosylvian gyrus showed the most intense labelling (i.e., the highest density of labelled soma across layers), with reactivity diminishing in more ventral regions. No labelled neurons were observed in layers I or IV across regions. This is consistent with the fact that SMI-32 clearly labels the somata and dendrites of pyramidal neurons (Del Rio and DeFelipe, 1994) while layers I and IV of sensory cortices typically contain primarily neuropils and stellate neurons, respectively. Conversely, the greatest immunoreactivity was consistently observed in layers III and V. Layer V and to a lesser extent VI contain large numbers of pyramidal neurons whose axons typically leave the cortex and project into the white matter. In contrast, layers II and III have smaller and fewer pyramidal neurons, whose connections are primarily corticocortical. Finally, all borders were easily identifiable at low magnification, and were “clear-cut” according to the criteria established by Mellott and colleagues (2010).

Overall, the patterns of SMI-32 reactivity observed in feline auditory cortex across the present study and that of Mellott and colleagues (2010) are in line with anatomical delineations based on other structural features. For example, regions that had higher somatic and dendritic counts based on SMI-32 labelling have also been shown to have higher myelin density (Robertson et al. 2024). This makes sense, as regions containing a greater number of axons are also likely to contain a greater number of myelinated fibers. The accordance between these two data sets suggests that regions of auditory cortex might be successfully discriminated based on a number of structural features (e.g., somatic density, myelin density) based on the nature of the available data.

## 4.1 Conclusion

The current study provides detailed quantifications and qualitative descriptions of the three auditory cortical regions that lie along the posterior ectosylvian gyrus of the feline brain. The soma counts, soma size estimates, and dendrite counts provided here supplement previous work describing the other 10 regions of the cat auditory cortex (Mellott et al., 2010). Together, these detailed quantifications provide insight into the anatomical composition of auditory cortex, and allow researchers to delineate auditory regions in a standardized way, allowing for interpretation and integration of results within a common spatial context.

## Acknowledgements

The authors would like to thank Katelyn Kittel for her contributions to animal care.

